# Systematic evaluation of high level visual deficits and lesions in posterior cerebral artery stroke

**DOI:** 10.1101/2022.05.19.492639

**Authors:** Ro Julia Robotham, Grace E Rice, Alex P Leff, Matthew A Lambon Ralph, Randi Starrfelt

## Abstract

Knowledge about the consequences of stroke on high level vision comes primarily from single case studies of patients selected based on their behavioural profiles with deficits in the recognition of a specific visual category such as faces or words. There are, however, no systematic, detailed, large-scale evaluations of the more typical clinical behavioural and lesion profiles of impairments in high level vision that may follow posterior cerebral artery (PCA) stroke. These goals were met by the current study through the data collected in the Back of the Brain (BoB) project: to date, the largest (N=64) and most detailed examination of patients with cortical PCA strokes selected based on lesion location rather than behavioural symptoms.

We present here two complementary analyses of the structural neuroimaging data and key indices of behavioural performance with the visual processing words, objects and faces: (1) a multivariate multiple regression analysis to establish the relationships between lesion volume, lesion laterality or the presence of a bilateral lesion with performance on words, objects and faces; and, (2) a voxel-based correlational method (VBCM) analysis to establish whether there are distinct or separate regions within the PCA territory that underpin the visual processing of these categories.

In contrast to the characterization of specific stroke syndromes like pure alexia or prosopagnosia in the literature, most patients in our cohort showed more general deficits in high level vision (n=22) or no deficits at all (n=21). Category-selective deficits were rare (n=6), and were only found for words, which, interestingly could follow left or right hemisphere lesions. The lesion analyses mainly confirmed the pattern reported in more selective cases: word recognition impairments are associated with a left-sided pattern of damage and face recognition deficits with a bilateral albeit right-dominant lesion pattern. Importantly, however, both general and more selective impairment may follow from left or right unilateral as well as bilateral lesions.

While the findings provide partial support for the relative laterality of posterior brain regions supporting reading in the left and, to a lesser extent, face processing in the right hemisphere, the results suggest that both hemispheres are involved in the visual processing of faces, words and objects. This has ramifications for researchers studying the healthy brain and for clinicians working with patients with PCA stroke. Clinicians are recommended to carry out formal assessment of face, word and object recognition as most patients are expected to present with a mixed picture of deficits.

## Introduction

Regarding the cerebral cortex, the posterior cerebral artery (PCA) supplies the occipital and ventral temporal lobes, regions involved in multi-level processing that leads to the identification of visual stimuli. Stroke within this territory (∼10% of stroke cases,^1^ often results in low-level visual deficits including hemianopia, but also higher-level visual deficits affecting the recognition of more complex stimuli such as faces, words and objects.^2-4^

While the consequences of stroke on low-level vision and ocular mobility have been investigated in large samples,^5,6^ there is a lack of systematic large-scale investigations of the clinical consequences of PCA stroke on high-level vision. Rather, knowledge about such consequences comes primarily from single case studies of patients with a selective or disproportionate deficit in the recognition of a specific category, who are typically selected based on their behavioural profiles. The most commonly described examples of selective higher-level visual deficits following PCA-stroke are in reading and face recognition. Single case studies suggest that pure prosopagnosia, a selective face recognition deficit, is typically caused by right hemisphere or bilateral lesions in the lateral mid fusiform gyrus,^7-11^ and that pure alexia, a selective reading deficit, is typically caused by lesions in the left posterior occipitotemporal gyrus or lateral mid fusiform gyrus.^12-16^ Functional neuroimaging studies of healthy participants have identified similar regions suggested to be category selective, namely the fusiform face area (FFA) and the visual word form area (VWFA).^17-19^ Recently, however, the focus has shifted from these core, category selective regions to the characterisation of the bilateral networks underlying high level vision, with various patient studies suggesting that the relationship between visual recognition deficits and lesion lateralisation might be less straightforward than previously assumed. ^20-23^

Here, we report data from the Back of the Brain (BoB) project, a systematic, detailed neuroimaging and neuropsychological examination of 64 patients with PCA stroke. The dataset is unique for two reasons: First, patients were recruited based on lesion localization within the cortical PCA-territory rather than behavioural symptoms. The study therefore gives us insights into the range and patterns of deficits that can be seen following PCA stroke and not just the rare patterns that are interesting enough to warrant a single case study. Secondly, high-level visual processing was assessed with a range of carefully matched tests of face, word and object stimuli, enabling direct comparison of performance across domains. Using these unrivalled data from the BoB project, we report here the variety of clinical presentations that follow PCA stroke, which in most cases is not a selective deficit, but rather is characterised by more general impairments in visual perception and recognition. In addition, we use brain-behaviour correlational methods to understand the types of lesions that cause impairment in visual recognition of faces, objects, and words, and which do not. We present two complementary analyses of the structural neuroimaging data and key indices of behavioural performance with words, objects, and faces: (1) a multivariate multiple regression analysis to establish the relationships between lesion volume, lesion laterality or the presence of a bilateral lesion with performance on words, objects and faces; and, (2) a voxel-based correlational method (VBCM) analysis to establish whether there are distinct or separate regions within the PCA territory that underpin the visual recognition of faces, objects and words.

## Materials and methods

### Participants

64 patients with a single stroke affecting cortex in the PCA territory (ischemic or haemorrhagic) occurring at least 9 months prior to participation, were recruited from two UK centres (University College London, University of Manchester) over a 24-month period. At the London site, patients were recruited from the PLORAS database^24^ and a specialist hemianopia clinic at the National Hospital for Neurology and Neurosurgery, University College London Hospitals, run by APL. At the Manchester site, patients were recruited from local clinics at Manchester Royal Infirmary, Salford Royal Hospital, The Walton Centre in Liverpool, and various speech therapy clinics in the Northwest region. To be included, patients had to have lesions affecting the cortical territory of the PCA; patients with lesions restricted to the brainstem, cerebellum, midbrain, and thalamus were excluded. Patients with bilateral infarcts were included as long as it was highly likely that they had suffered a single episode of stroke. Patients with head injuries, or diagnosed developmental, psychiatric, or other neurological disorders were excluded. 46 age-matched control participants were also included.^25^

Table 1 summarises the demographic information and background neuropsychological data. All participants were native English speakers, and mainly right handed. The laterality subgroups were not selected to be matched across demographic variables, but were not significantly different in terms of age, education level or time since stroke (all p’ s>0.1, see supplementary Table 1 for details). All patients underwent visual field (a.m. Nordfang et al.^26^ and visual acuity testing.^27^ Visual field defects were found in 60 patients (94%). 26 (41%) patients had homonymous hemianopia and 28 (44%) had quadrantanopia (lower = 13, upper = 15). Six patients had bilateral visual field deficits (9%). All patients had normal or corrected to normal visual acuity.

**Table 1:** Participant demographics and background neuropsychological testing. All values represent averages, with standard deviation in parentheses, with the exception of gender and handedness which represent counts.

| Demographics | Control Total | Patient Total | Left | Bilateral | Right |
| --- | --- | --- | --- | --- | --- |
| N | 46 | 64 | 32 | 9 | 23 |
| Age | 61.5 (14.6) | 60.9 (13.1) | 63.9 (11.6) | 57.6 (10.7) | 57.9 (15.2) |
| Gender (M/F) | 22/24 | 52/12 | 26/6 | 8/1 | 18/5 |
| Education (years) | 15.2 (1.9) | 14.0 (2.7) | 14.0 (2.5) | 13.8 (3.6) | 14.3 (2.6) |
| Handedness (LH/Mixed/RH) | 2/2/42 | 6/1/57 | 5/1/26 | 1/0/8 | 0/0/23 |
| Time since stroke (months) |  | 41.9 (49.7) | 42.3 (48.0) | 40.0 (28.5) | 42.0 (59.4) |
| Lesion volume (cm<sup>3</sup>) |  | 37.0 (35.5) | 31.8 (29.9) | 61.4 (37.9) | 34.7 (39.2) |
| Background neuropsychology |  |  |  |  |  |
| Geriatric Depression Scale <sup>28</sup><br>(max 15) |  | 3.68 (3.54) | 3.41 (3.13) | 5.00 (4.95) | 3.52 (3.50) |
| Oxford Cognitive Screen <sup>29</sup><br>(Impaired tests - max 10) |  | 0.92 (1.29) | 0.84 (1.25) | 1.44 (1.94) | 0.83 (1.03) |
| WAIS-IV <sup>30</sup> Digit Span<br>(Forward, max = 16) | 10.83 (2.15) | 10.09 (2.24) | 9.94 (2.18) | 11.33 (2.78) | 9.83 (2.04) |
| WAIS-IV <sup>30</sup> Digit Span<br>(Backward, max = 14) | 7.55 (2.12) | 6.47 (2.10) | 6.28 (2.16) | 6.33 (1.66) | 6.78 (2.21) |

Legend: WAIS-IV: Wechsler Adult Intelligence Scale IV

### Structural scanning

Structural brain imaging data were acquired in all patients and 22 of the control participants. Structural scans were acquired on two 3T Phillips Achieva scanners with 32-channel head-coils and a SENSE factor of 2.5 in London and Manchester. A high-resolution T1 weighted structural scan was acquired including 260 slices covering the whole brain with TR = 8.4ms, TE = 3.9ms, flip angle = 8 degrees, FOV = 240 × 191mm^2^, resolution matrix = 256 × 206, voxels size = 0.9 × 1.7 × 0.9mm^3^.

### Automated lesion identification procedure

Automated outlines of the area affected by stroke were generated using Seghier *et al*.^31^’ s modified segmentation-normalisation procedure (run using SPM12), which is designed for use with brain-injured patients and identifies areas of lesioned tissue in order to optimise fitting lesioned brains to standard MNI space. Segmented images were smoothed with an 8mm full-width half maximum Gaussian kernel and submitted to the automated lesion identification and definition modules using the default parameters. The automated method involves initial segmentation and normalising into grey matter, white matter, CSF, and an extra tissue class for the presence of a lesion. After smoothing, voxels that emerge as outliers relative to the control participants’ scans are identified and the union of these outliers generates the “fuzzy lesion map” from which the lesion outline is derived. Using this procedure, there were four patients whose small lesions could not be identified. For these patients, a neurologist (APL) manually traced the lesions using a semi-structured lesion identification technique, using the fuzzy lesion map to guide tracing. The “fuzzy lesion” image was used to calculate the lesion variance image (Fig. 1B) and as input to the VBCM analysis (Fig. 2). The binarised lesion image was used to create the lesion overlap map in Fig. 1A.

**Figure 1:**
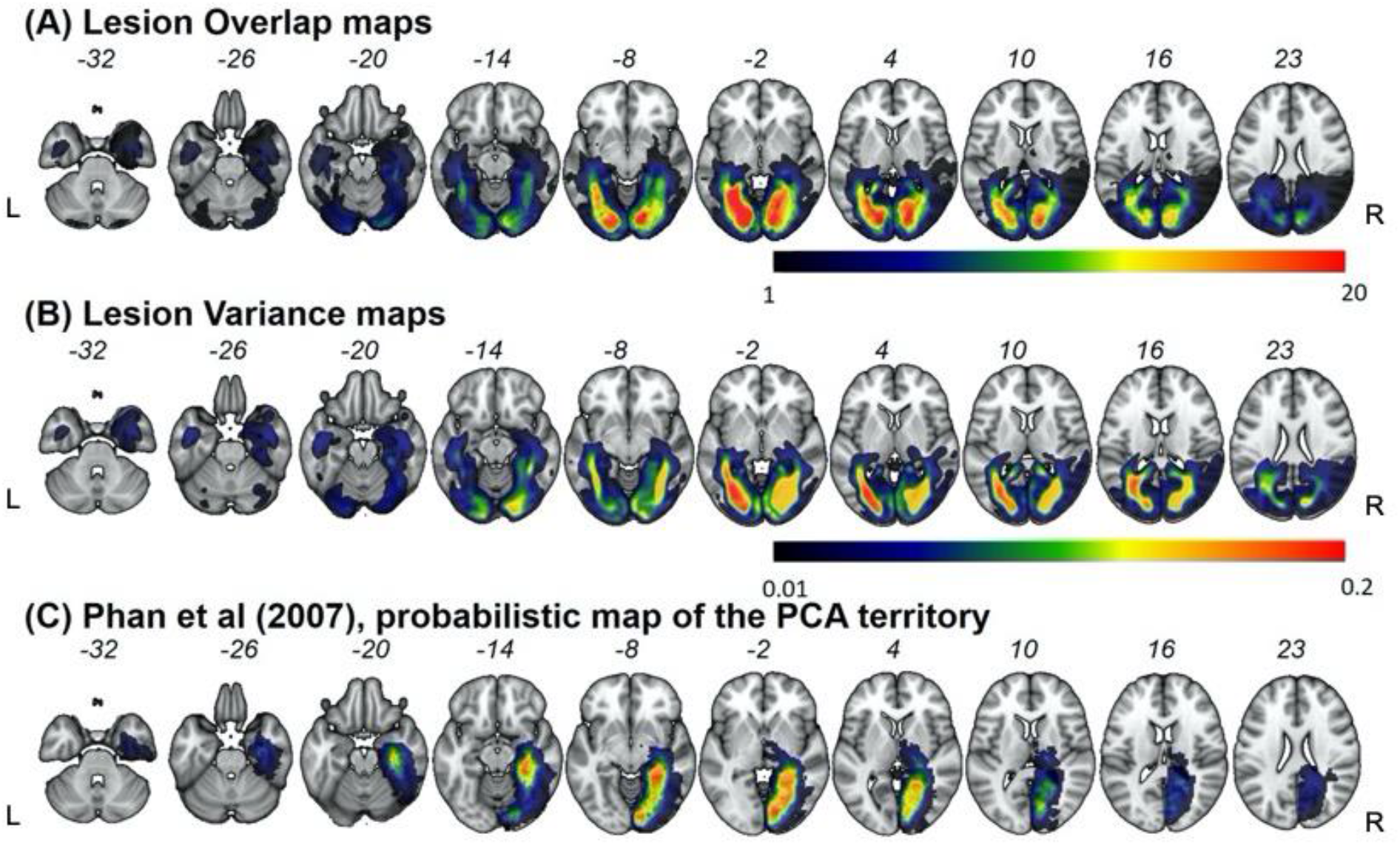
Lesion overlap and variance maps for the 64 PCA stroke cases. L= left hemisphere, R = Right hemisphere. (A) Lesion overlap defined by the method described in Seghier *et al*.^31^. Colour bar indicates the number of patients with lesion in that area. Warmer colours = greater overlap, cooler colours = less overlap. (B) Lesion variance map. Colour bar indicates the variation at each voxel across the PCA territory. Warmer colours = greater variability, cooler colours = less variability. (C) Probabilistic definition of the PCA territory (reproduced with permission from Phan *et al*.^42^).

**Figure 2:**
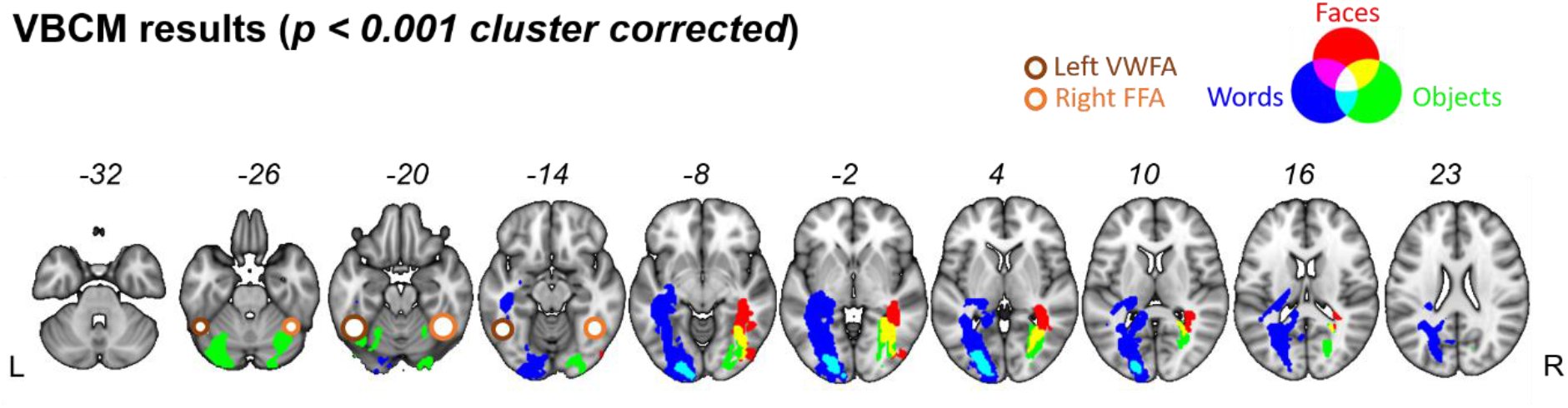
VBCM results of structural correlates of Word, Object and Face Recognition (*p < 0*.*001 cluster corrected*). Results from the three domains are overlaid on one another (word recognition, blue; object recognition, green; face recognition, red). Overlap between words and object recognition are shown in cyan. Overlap between objects and faces are shown in yellow. All clusters were obtained by applying a voxel-level threshold of p < 0.001, and a family-wise cluster correction of p < 0.05. Left visual word form area (Jobard et al., 2003) and right fusiform face area (Muller et al., 2018) illustrated with brown circles.

### Behavioural Assessment

All patients completed a detailed neuropsychological battery, which was designed to test, systematically, a broad range of visual perceptual functions (described in detail in Robotham *et al*.^32^). Here, we report results from tests of face, word and object recognition. A central aim of the project was to assess word, object, and face recognition abilities in comparable ways in order to assess both the range and specificity of visual perceptual deficits following PCA-stroke. Commonly, tests of word, object, and face recognition have different task demands (e.g., naming vs forced choice), and /or they tap different levels of processing (e.g., visual perception vs. memory), leaving it unclear if reported dissociations are between stimulus categories, types of processing, or task demands. Devising tasks that test face and word recognition in comparable ways has been a particular challenge for the field. ^32,33^ To overcome this potential pitfall, we designed and selected tests that were comparable in experimental setup and response mode across domains, including tests of delayed matching, recognition memory, familiarity judgements, and naming for each stimulus type.

### Experimental design

The various assessments are briefly described below. Full details are available in Robotham *et al*.^32^.

#### Delayed matching and surprise recognition test

This test was designed specifically for the BoB project. The first part was a *Delayed matching task*. First a stimulus was presented for 1000ms at the centre for the screen followed by a 1000ms blank screen. Then a probe was presented for 180ms at the centre of the screen. Participants were asked to decide if the probe was identical to the target or not. The probe was either larger or smaller than the target, to ensure that mere change detection was not sufficient to perform the task. Uncropped faces, lower case words and common objects were tested in separate blocks. 12 stimuli were used for each category. This was followed by a *Surprise recognition test* in which two stimuli were presented simultaneously vertically on the screen: one familiar and one novel stimulus. Participants were asked to determine which of the images they had seen before. The familiar stimuli consisted of the 36 stimuli used in the Delayed matching test and 36 novel stimuli (individually matched with a high degree of similarity to the familiar stimuli).

#### Familiarity judgements

Tasks requiring differentiation between familiar (words, objects, famous faces) and unfamiliar (nonwords, nonsense objects, unfamiliar faces) visual items were included for each domain to assess visual recognition without the need for naming out loud. For all domains, stimuli were presented centrally on a screen, and participants indicated via button-press, as quickly and accurately as possible, whether the item was familiar or not. The dependent variables were accuracy and correct RT. The Lexical decision task included 30 words and 30 pseudowords of either 3, 5, or 7 letters in length (selected from the task used by Behrmann & Plaut^21^). The Object decision test used 36 objects and 36 chimeric non-objects.^34,35^ The Face familiarity decision test contained the 40 famous faces included in the Famous Face Naming task (see below, the familiarity task was always presented first) and 40 unfamiliar faces. This test was designed specifically for the Back of the Brain project.

#### Naming

A naming test was included for each domain. Participants were asked to name stimuli as quickly and accurately as possible. Accuracy was recorded by the experimenter, and RT from stimulus onset to vocal response was measured (for words and objects, see below). For the *Reading* task, participants were asked to name out loud 75 regularly spelled single words of 3, 5, or 7 letters in length.^14,36^ Each word was displayed on the screen until a response was recorded or a maximum of 4 seconds. Responses provided after 4 seconds were scored as errors. In the *Picture Naming* task, participants were required to name 45 black and white line drawings of objects. The stimuli were 15 non-living items and 30 living items. Items were presented on the screen until a response was made or for a maximum of 6 seconds. Responses over 6 seconds were scored as errors. The *Famous Face Naming* task included 40 pictures of famous faces. Participants were asked to name the faces. If participants were unable to provide a name, recognition of the person was tested (e.g., provision of why the person is famous, what they do, where they live etc.). Only the naming score was included in the present context. The main measure for this test is accuracy, reaction time data were not scored due to the extensive verbal output.

### Analysis of behavioural results

To take advantage of the richness of data collected in these tests, while enabling direct comparison across domains, composite scores were calculated to provide a summary measure of performance for each of the three domains of interest (words, objects and faces). This composite score was generated by using unrotated fixed-factor principal components analysis to create a single weighted average of the combined accuracy and RT, for each patient and control participant. To assess the presence of a deficit in each domain, the performance of each individual patient on each composite score was compared to the control group using single case statistics.^37^

## Lesion analyses

### Multiple Regression Analysis (multivariate analysis)

First, we sought to establish the relationship between the patients’ lesions and their behavioural performance on word, object and face recognition. Specifically, we explored the effect of (a) total lesion volume, (b) lesion laterality (left vs. right), and (c) the effect of a unilateral vs. a bilateral lesion.

To quantify the lesions across the patient group, a mask of the PCA territory in the left and right hemisphere was derived from the Harvard Oxford atlas^38^ and the John Hopkins White Matter atlas^39^. The PCA mask consisted of the occipital pole, calcarine sulcus (inferior, superior, intracalcarine), lingual gyrus, parahippocampal gyrus (posterior, anterior), fusiform gyrus (occipital, temporal occipital, posterior, anterior), inferior temporal gyrus (temporal occipital, posterior), lateral occipital cortex (inferior, superior) and the precuneus. The white matter tracts of the inferior longitudinal fasciculus and splenium (including the forceps major) were also included.

The proportion of overlap between each patient’ s lesion and the left and right PCA mask was calculated. Each patient’ s lesion was defined as the binary lesion image from the automated lesion identification method above.^31^ The proportion of lesion overlap for each patient was then used to calculate three measures: (1) Total lesion volume (the sum of left + right PCA overlap), (2) Lesion laterality (the difference between left - right PCA overlap), and (3) the presence of a unilateral vs. bilateral lesion, coded as either 1 (unilateral lesion) or 2 (bilateral lesion). We included both lesion laterality and the presence of a unilateral vs. bilateral lesion in the models in order to differentiate between the effect of a large unilateral lesion affecting a critical functional area in one hemisphere, and the presence of a large bilateral lesion affecting the same functional area in both hemispheres.

To understand how lesion volume, lesion laterality and the presence of a bilateral lesion influenced visual perceptual performance, we built a linear regression model to test the relationship between the three lesion measures as independent variables and the performance on one of the composite scores (words, objects, faces). Separate simultaneous linear regression models were calculated for each domain and were run using SPSS (version 25).

In addition to examining the overlap between each patient’ s lesion and the left and right PCA masks as a whole, we also calculated lesion overlap within each constituent ROI within the PCA mask. We conducted further linear regression analyses to assess the importance of specific subregions of the PCA territory (listed above) in visual perceptual performance.

### Voxel-based correlational methodology (VBCM) analysis

VBCM was implemented to further explore which regions within the PCA territory were associated with visual perceptual performance.^40^ VBCM is a variant of voxel-based lesion symptom mapping,^41^ in which both the behaviour and signal intensity measures are treated as continuous variables. This analysis was conducted in SPM12 using the smoothed fuzzy lesion maps (which contain both the grey and white matter), where each voxel represents the % abnormality. For this analysis, we assumed a negative correlation between tissue abnormality and the behavioural composite score (i.e., greater abnormality leads to worse performance). The patients’ composite scores for the three domains (words, objects, faces) were entered into separate VBCM analyses, along with covariates of age (continuous variable) and site of scanning (London or Manchester: categorical variable) to account for intensity differences between scanners. In a separate analysis, total lesion volume in the left and right hemisphere were also included as additional covariates. Unless otherwise noted, a threshold at voxel-level p < 0.001 and family-wise error corrected (FWEc) cluster-level p < 0.05 was applied.

### Data availability

The data that support the findings of this study are available on request from the corresponding author. The data are not publicly available as they contain information that could compromise the privacy of research participants.

## Results

### Lesion profiles

Fig. 1A shows the lesion overlap map for all patients. Lesions covered the PCA territory and aligned with previous descriptions of PCA infarcts.^15,42^ Five of the nine bilateral cases showed more damage in the right hemisphere compared to the left; one patient showed more damage in the left hemisphere compared to the right; three patients showed no hemispheric differences in lesion volume. The bilateral group had larger lesions on average than the left hemisphere group (Table 1; t (39) = 2.48, p = 0.02). No other group differences were significant (left vs. right t (53) = 0.32, p = 0.75; right vs. bilateral: t (30) = 1.75, p = 0.09).

The maximal lesion overlap was in the medial occipital lobe, posterior lingual gyrus and medial posterior fusiform gyrus (Fig. 1A, red). Despite the homogenous overlap within the PCA territory, there were differing degrees of variability across the two hemispheres (Fig. 1B). Within the left hemisphere there was a greater degree of variability, compared to the right hemisphere. This relative lack of variability in the right hemisphere was caused by a number of right hemisphere patients with large (and similar) lesions. Notably, there was limited extension into the lateral aspects of the posterior fusiform gyrus (purported by Fmri explorations in healthy participants to be the critical region for category-specific responses in both hemispheres). Two patients showed a degree of overlap with the left hemisphere VWFA (1 left hemisphere, 1 bilateral; coordinates defined from Jobard *et al*.^43^), and nine patients showed a degree of overlap with the right hemisphere FFA (6 RH, 3 bilateral; coordinates defined from Muller *et al*.^44^). Critically, no patient in the BoB cohort had an isolated lesion affecting only the lateral posterior fusiform gyrus, posited to be the core site of the FFA.

### Behavioural profiles

Behavioural performance in word, object and face recognition was assessed using composite scores. The factor loadings of the individual measures on the composites are available in Supplementary Table 2 (see also Rice *et al*.^25^). Composite scores for individual patients and controls are available in the Supplementary Table 3. There were high and significant correlations between the composite scores for both patients and controls (Patients: Words-Objects: *r* = .812 (95% CI [.707, .882]; Words-Faces: *r* = .679 (95% CI [.520, .793],; Faces-Objects: *r* = .824 (95% CI [.725, .889],; Controls: Words-Objects: *r* = .687 (95% CI [.496, .825]); Words-Faces: *r* = .478 (95% CI [.218, .675]; Faces-Objects: *r* = .613 (95% CI [.393, .767], all *p* < .001). Deficits within each domain for each patient were determined using single case statistics^37^ (Table 2). One third of the patients were significantly impaired in all three domains, and this occurred following left and right unilateral as well as bilateral lesions. Another third of the patient group showed no significant deficit in either domain. The remaining third showed more selective deficits. Most of these showed deficits in two domains (n = 14) while a few showed deficits confined to one category. Three patients with left hemisphere lesions and three right-handed patients with right hemisphere lesions showed a selective deficits for words. No patients showed a selective deficit for faces or objects.

**Table 2.** Patterns of deficits across domains. Significant deficits were determined by comparing each patient’ s scores to the control group using the Bayesian test for a deficit allowing for covariates.^37^.

|  | Left hemisphere<br>(n=32) | Bilateral<br>(n=9) | Right hemisphere (n=23) | Total<br>(N=64) |
| --- | --- | --- | --- | --- |
| **No deficits** | 12 | 1 | 8 | 21 |
| **WOF** | 10 | 6 | 6 | 22 |
| **WF** | 2 | 0 | 1 | 3 |
| **WO** | 5 | 0 | 4 | 9 |
| **FO** | 0 | 2 | 1 | 3 |
| **W only** | 3 | 0 | 3 | 6 |
| **O only** | 0 | 0 | 0 | 0 |
| **F only** | 0 | 0 | 0 | 0 |

Legend: W=word deficit, F=face deficit, O=object deficit

**Table 3:**
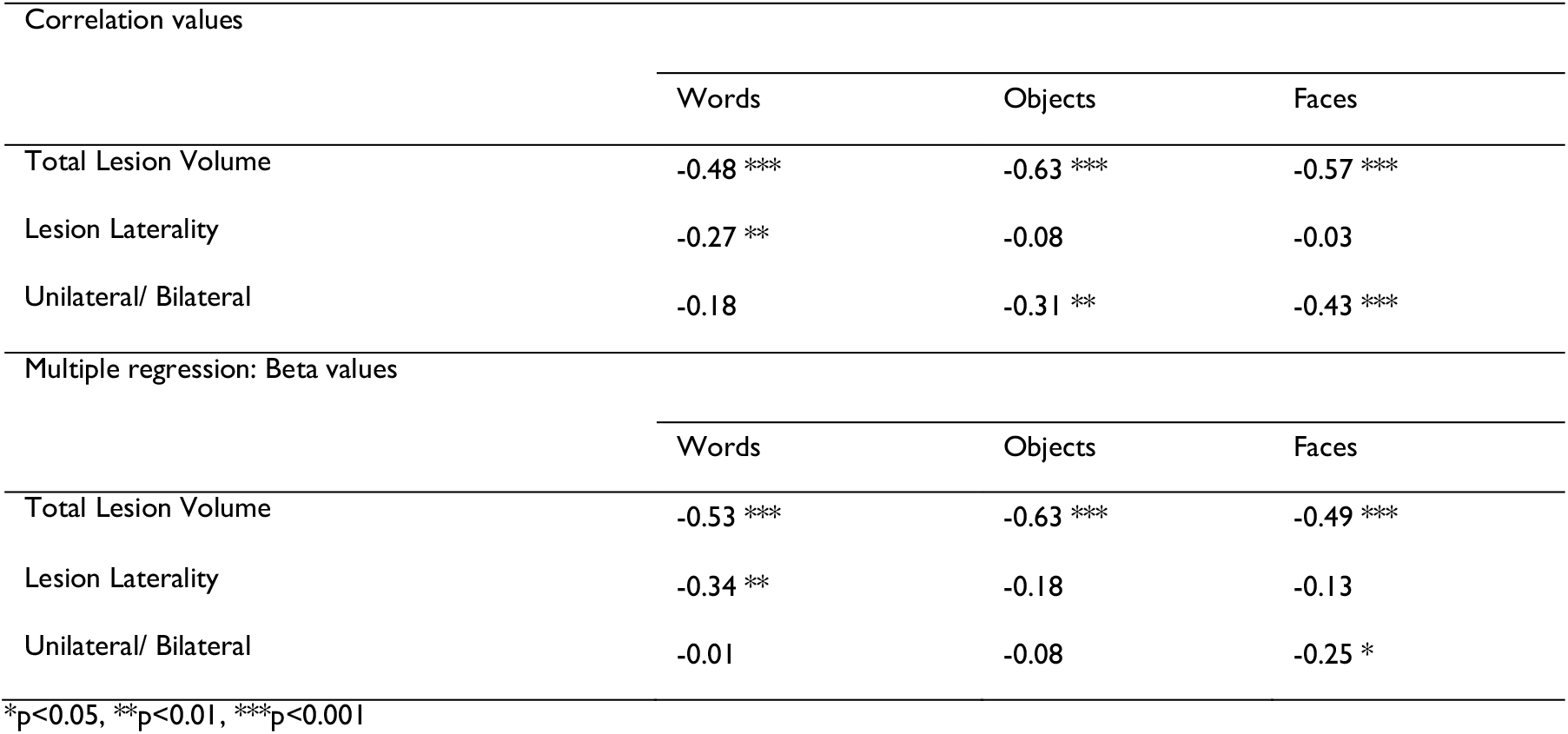
Whole brain multiple regression results.

### Multiple regression (whole brain analysis)

The multiple regression analyses (Table 3) of lesion profile (total lesion volume, lesion laterality, the presence of a bilateral lesion) and level of impairments in visual processing indexed by composite scores showed that total lesion volume was most strongly related to performance (words: beta = −0.53, t (63) = 4.60, p = p < 0.0001; objects: beta = −0.63, t (63) = 5.88, p < 0.0001; faces: beta = −0.48, t (63) = 4.38, p < 0.0001). This was the only lesion factor that was significantly related to performance on the object composite score. Performance on the word composite score was additionally related to lesion laterality (beta = −0.34, t (63) = 3.25, p = 0.002), driven by poorer word recognition performance following a left hemisphere lesion. Poorer performance on the face composite score was related to the presence of a bilateral lesion (beta = −0.25, t (63) = 2.26, p = 0.03).

The same analysis was conducted on the constituent ROIs within the PCA mask. In this analysis, the three PCA lesion measures were included in a step-wise regression alongside the proportion of damage in each constituent PCA ROI (see Supplementary Table 4 for full results). Thus this analysis tested whether damage to a specific subregion explained performance over and above total lesion volume or laterality. Aligning with the previous results, performance on the word composite score was significantly related to lesions of predominantly left hemisphere ROIs (left ILF: beta = −0.49, t (63) = 5.45, p < 0.0001; left occipital lobe: beta = −0.39, t (63) = 4.28, p < 0.0001), and right pITG (beta = −0.22, t (63) = 2.62, p = 0.011), the latter aligning with the behavioural observation of word deficits in the right hemisphere patients. Performance on the object composite score was significantly related to total lesion volume (beta = −0.57, t (63) = 5.77, p < 0.0001), and left ILF damage (beta = - 0.23, t (63) = 2.53, p = 0.02). Finally, performance on the face composite score was related to total lesion volume (beta = −0.61, t (63) = 4.54, p < 0.0001), the presence of a bilateral lesion (beta = 0.27, t (63) = 2.53, p = 0.014), and also to damage to the right aITG (beta = −0.21, t (63) = 2.09, p = 0.04) and right lingual gyrus (beta = −0.29, t (63) = 2.19, p = 0.03)

## VBCM analysis

As well as exploring brain-behaviour mapping in the core PCA ROI regions, we used VBCM to provide a whole-brain analysis (Fig. 2). The results replicated those found in the ROI-based analysis, and reinforced the patterns shown in the behavioural analysis of these data. Performance on the word composite score correlated with a left hemisphere cluster extending from the occipital pole, along the fusiform and lingual gyri (Fig. 2; blue). This cluster also encompassed the white matter of the ILF and splenium/forceps major, which have long been hypothesised to play a role in pure alexia ^45,46^. Interestingly, the word cluster remained exclusively within the left hemisphere even at a lower threshold (P<0.01) (Fig. 3). Performance on the object composite score correlated with bilateral clusters in the left occipital pole, and the right ILF (Fig. 2; green). The cluster in the left occipital lobe overlapped with the word cluster (Fig. 2; cyan). Finally, performance on the face composite score correlated with a right hemisphere cluster mainly within the white matter of the IFOF and ILF. This cluster overlapped with the object cluster in the right ILF (Fig. 2; yellow). At the lower threshold, significant clusters for faces were also revealed in the left hemisphere (Fig. 2). The ROI based regression analyses showed that total lesion volume was the most strongly related to behavioural performance. This finding was replicated in the VBCM analysis, as inclusion of lesion volume (calculated for the left and right hemisphere separately) removed most of the significant clusters, particularly for the object and face composite scores.

**Figure 3:**
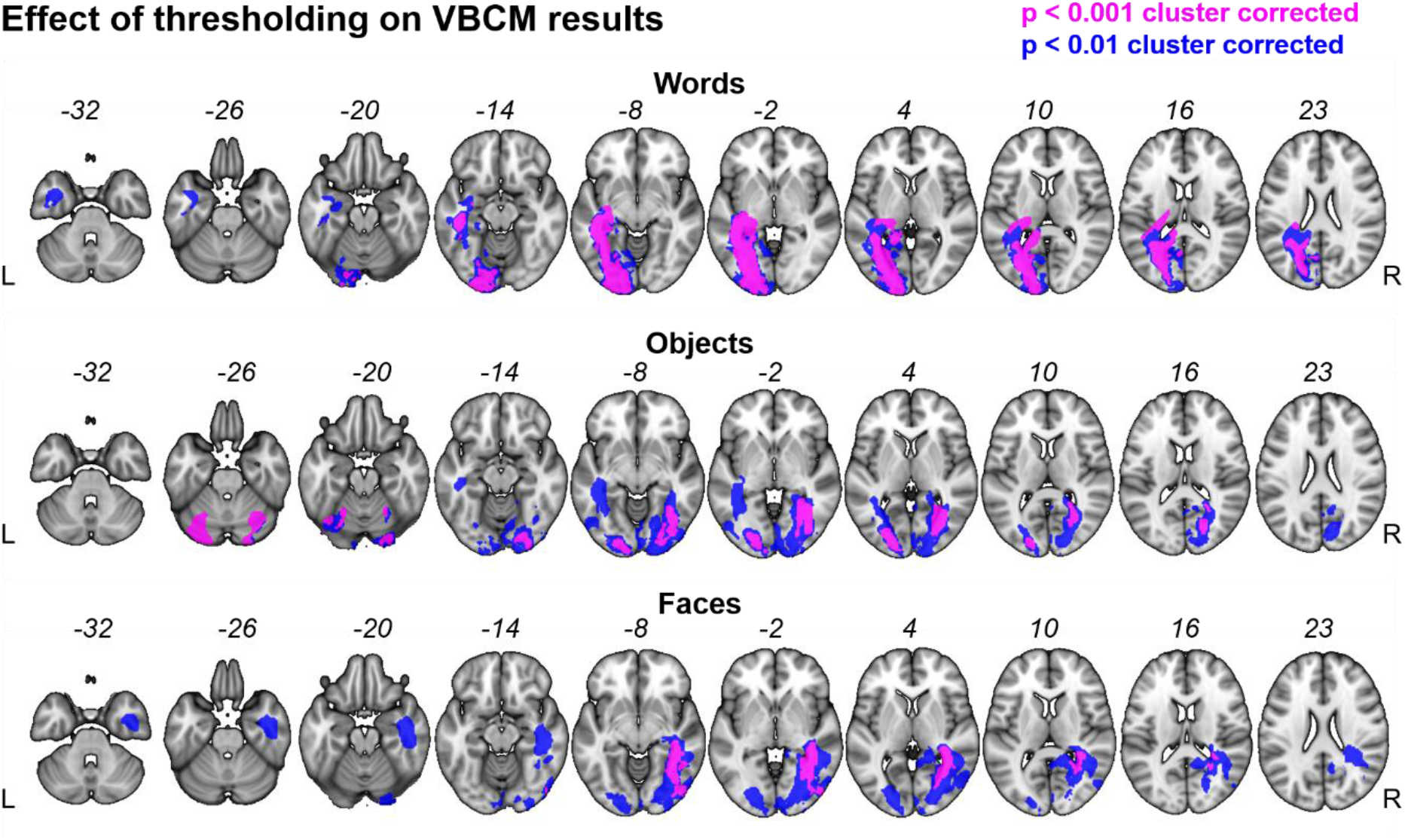
Clusters in blue were obtained by applying a more liberal voxel-level threshold of p < 0.01, and a family-wise cluster correction of p < 0.05. Overlap with high-threshold clusters with p < 0.001 (shown in figure 2) are shown in purple.

## Discussion

Damage to the brain regions supplied by the posterior cerebral artery characteristically results in visual perceptual deficits, but most of our knowledge about such deficits come from single case studies or smaller case series of patients selected based on selective or disproportionate deficits in the recognition of specific visual categories.^12^–^14^,^47^–^49^ Here, we adopted an alternative contemporary approach that is of more use to researchers and clinicians: completing a large-scale, in-depth systematic evaluation to map the relationship between lesion location following PCA territory stroke and high-level visual perceptual performance with words, objects and faces in a large group of patients selected based on lesion location rather than cognitive profile. The BoB project contains representative coverage of the entire PCA territory,^42^ with significant variability of lesion size within both hemispheres and therefore provides novel insights into the diversity of visual perceptual profiles that can arise following PCA stroke.

Behaviourally, the key findings were that 1) very few patients showed selective deficits in only one domain, 2) about one third of the patients showed significant impairment across domains, and this could follow unilateral lesions to either hemisphere as well as bilateral lesions, and 3) about a third of the patients performed within the normal range across all three domains, thus constituting a central comparison group showing that general slowness or nonspecific effects of PCA stroke are not sufficient to impair performance on the behavioural measures applied. The observed deficits in the remaining patients can therefore be considered to result from their specific lesions rather than general effects of having suffered a stroke. Only six patients out of the 64 included showed a selective deficit, and this was only observed for words, suggesting that selective deficits are indeed rare. In most cases, reading deficits occurred together with object recognition deficits. The same was the case for face recognition deficits.

Linking behaviour to lesions, we found that across all three domains, total lesion volume had the strongest relationship with behavioural performance. Aligning with results from the literature on cases selected based on behavioural performance/impairment,^12,47^ word recognition performance was also related to lesion laterality; patients with left hemisphere lesions performed worse with written words. Face recognition performance, however, was not related to lesion laterality but instead to lesion volume across hemispheres. Overall, results from ROI-based multiple regression and the VBCM analysis showed the same pattern of results: In both analyses we found that across all three domains, total lesion volume had the strongest relationship with behavioural performance.

In the VBCM analysis the majority of the significant clusters were found within the white matter (particularly within the territory of the inferior longitudinal fasiculus). This seems to align with the classical hypothesis that disconnection of a functional region may give rise to the same behavioural deficit as direct damage in pure alexia and prosopagnosia. ^7^,^50^–^54^

Regarding lesion lateralisation, the findings provide a nuanced picture: the lesion analyses align well with the literature on more selective deficits, with left hemisphere regions being associated with word performance, and a bilateral but right dominant set of regions were associated with face recognition impairment. In addition, it is also clear that impairment in either category might follow from lesions to either hemisphere. The extent of lateralisation of face and word recognition has been highly debated both within the patient and neuroimaging literature.^20,21,55,56^ Examples of patients from the single case literature have been used to argue that face and word processing rely on largely lateralised and relatively independent cognitive processes. While almost all patients with pure alexia have left hemisphere lesions,^12,57^ patients with pure prosopagnosia typically either have bilateral lesions, or lesions in the right hemisphere.^58,59^ Early studies using fMRI provided additional evidence that face and word processing were highly lateralised: A region in the left occipitotemporal gyrus (VWFA) was shown to be more responsive to words than low level stimuli and consonant strings ^17,19^ and a region in the right occipitotemporal gyrus (FFA), was shown to be more responsive to faces than scrambled faces.^60,61^ However, contemporary more sensitive fMRI has shown that neither faces nor words lead to fully lateralised activation. Both categories generate bilateral activation with varying degrees of asymmetry with words leading to a stronger left lateralised response than the right lateralised response that faces give rise to.^62-64^ There are also rare examples of patients with prosopagnosia following a left hemisphere lesion and pure alexia following a right hemisphere lesion suggesting that both hemispheres provide substantial contributions to face and word recognition,^65-68^ and there is increasing evidence that face and word recognition impairment is typically associated rather than dissociated following brain injury.^21,22,69^

In the current study, patients with left hemisphere lesions did perform worse as a group with written words suggesting a left hemisphere dominance for words. In the VBCM-analysis, performance on the word composite score correlated with a left hemisphere cluster extending from the occipital pole, along the fusiform and lingual gyri, and encompassing the white matter of the ILF and splenium/forceps major. This aligns well with the literature on pure alexia, where damage or disconnection of the left mid fusiform gyrus in particular has been suggested to be critical.^12,47,70^ There were however also patients with lesions restricted to the right hemisphere who had a poor visual word processing performance. In fact, three of the six patients in our sample who had a selective deficit in word recognition (with preserved object and face recognition) had lesions restricted to the right hemisphere. None of these patients were left handed. While their word recognition deficit was milder than the deficit measured in three patients with lesions in the left hemisphere, the findings suggest that the right hemisphere also provides important contributions to word recognition. In line with this, the ROI-analysis pointed to bilateral contributions to word recognition. Poor word processing was significantly related not only to ROIs in left hemisphere, but also to the right pITG, a region that has been implicated in neglect dyslexia.^71,72^ Taken together, our findings regarding the cerebral substrates of visual word recognition supplement the existing literature and provide additional evidence that while visual word recognition is strongly lateralised to the left hemisphere, regions in the right hemisphere also makes critical contributions.

Face recognition, though considered to be more bilaterally distributed than word recognition, is still thought to be somewhat lateralised to the right.^58,60,73^ In our sample, no patient showed a selective deficit for faces, but face processing problems were observed following unilateral lesions to either the right or left hemisphere, and also following bilateral lesions. The VBCM analysis did however reveal that face processing correlated with a cluster in the right hemisphere, mainly within the Inferior fronto-occipital fasciculus (IFOF) and the Inferior longitudinal fasciculus (ILF). At the lower threshold significant clusters in the left hemisphere were also revealed for faces. Taken together, our results provide additional evidence that the neural correlates for face recognition are highly bilaterally distributed, but with some degree of lateralisation to the right, and additional evidence that face recognition likely relies on the integrity of both the ILF and the IFOF. The ILF connects the occipital and temporal-occipital areas to anterior temporal areas and is therefore strategically placed in relation to the occipital face area (located in the inferior occipital gyrus) and the fusiform face area (in the posterior and middle fusiform gyrus) that are considered key regions of the core face network.^74,75^ The IFOF also begins in the ventral occipital lobe but terminates in the frontal cortex and, the inferior frontal gyrus is considered as part of the extended face network. The IFOF is also thought to play a role for face recognition, maybe more specifically related to remembering faces. Differences in face processing abilities have been related to integrity of the ILF and the IFOF ^76^ and patients with congenital prosopagnosia have been shown to have a reduction in structural integrity of both tracts bilaterally.^77^

No patient showed a selective deficit for objects, which is not surprising given that isolated visual object agnosia without alexia or prosopagnosia is not thought to occur.^78,79^ Including the object category in the assessments was important, however, to determine the possible selectivity of face and word recognition deficits in the cohort. Had we only compared word and face processing, the observed pattern would be very different and more indicative of a category selective organization. The object composite score was only related to lesion volume and not to lesion laterality or the presence of a bilateral lesion. According to the VBCM analysis, poor performance on object composite score was related to damage to the left ILF. This is interesting as object recognition has not traditionally been considered to rely on lateralised processes. While there are reports of object agnosia following unilateral lesions to the right or left hemisphere,^80-84^ these cases typically also have either prosopagnosia or alexia, depending on the hemisphere. The majority of visual agnosia cases, however, have bilateral lesions of ventral occipitotemporal cortex,^79^ and even in unilateral cases, functional imaging has demonstrated abnormal activation patterns also in the contralesional hemisphere.^81,83,84^

In conclusion, while our findings offer partial support for the relative laterality of posterior brain regions supporting reading (left) and, to a lesser extent, face processing (right); there are two important caveats. Firstly, for all three categories, there is clear evidence that both hemispheres are involved in higher-order processing; this has ramifications for those studying processing in the undamaged brain (e.g., functional neuroimagers) and those interested in rehabilitating patients with visual perceptual disorders. Secondly, these results will help guide clinicians in what to expect in terms of higher-order visual deficits in the next patient they see with PCA stroke: it’ s likely to be a mixed picture, so we suggest formal assessment of reading, face and object perception in all cases.

## Acknowledgements

We would like to thank the patients and their families for their generosity of time and patience during their participation in the project. We would also like to thank Sheila K. Kerry for her contributions to recruitment, data collection and preliminary analyses, Nicolaj Mistarz for help with coding of the behavioural data, and the Friends of Fakutsi Association (FFA) for support during project development.

## Supplementary material

Supplementary material is available at *Brain* online.

