## Supplementary material for "Systematic evaluation of high level visual deficits and lesions in posterior cerebral artery stroke"

**Supplementary Table 1. Comparison of demographics for laterality subgroups.**

| Demographics | Left | Bilateral | Right |
| --- | --- | --- | --- |
| N | 32 | 9 | 23 |
| Age | 63.9 (11.6) | 57.6 (10.7) | 57.9 (15.2) |
| Education (years) | 14.0 (2.5) | 13.8 (3.6) | 14.3 (2.6) |
| Time since stroke (months) | 42.3 (48.0) | 40.0 (28.5) | 42.0 (59.4) |
| **Comparison of demographics for laterality groups** | | | |
|  | Left vs Right  (*df* = 53) | Left vs Bilateral (*df* = 39) | Right vs Bilateral (*df* = 30) |
| Age | *t* = 1.69, *p* = 0.10 | *t* = 1.48, *p* = 0.15 | *t* = 0.06, *p* = 0.95 |
| Education (years) | *t* = 0.42, *p* = 0.68 | *t* = 0.18, *p* = 0.86 | *t* = 0.42, *p* = 0.68 |
| Time since stroke (months) | *t* = 0.02, *p* = 0.98 | *t* = 0.14, *p* = 0.89 | *t* = 0.09, *p* = 0.92 |

**Supplementary Table 2. Component loading tables for the unrotated PCA composite scores.** Any value above 0.5 is considered to be significant – all tests load significantly on their respective composite scores with the exception of the RT in the picture naming task (which falls just below)

**Word composite**

| **Test** | **Component Loading** |
| --- | --- |
| Word Reading (Accuracy) | **0.916** |
| Word Reading (3 letter words RT) | **0.908** |
| Lexical Decision (Real word RT) | **0.873** |
| Word Delayed Matching (Accuracy) | **0.870** |
| Word Surprise Recognition (RT) | **0.865** |
| Lexical Decision (Accuracy) | **0.858** |
| Word Delayed Matching (RT) | **0.814** |
| Word Surprise Recognition (Accuracy) | **0.514** |

**Object composite**

| **Test** | **Component Loading** |
| --- | --- |
| Object Decision (Accuracy) | **0.854** |
| Object Delayed Matching (RT) | **0.799** |
| Object Surprise Recognition (RT) | **0.793** |
| Picture Naming (Accuracy) | **0.775** |
| Object Delayed Matching (Accuracy) | **0.754** |
| Object Decision (Real Object RT) | **0.738** |
| Object Surprise Recognition (Accuracy) | **0.688** |
| Picture Naming (RT) | **0.401** |

**Face composite**

| **Test** | **Component Loading** |
| --- | --- |
| Famous Face Recognition (Accuracy) | **0.894** |
| Famous Face Naming (Accuracy) | **0.849** |
| Face Delayed Matching (Accuracy) | **0.841** |
| Face Surprise Recognition (Accuracy) | **0.822** |
| Face Familiarity (Accuracy) | **0.810** |
| Face Familiarity (Familiar RT) | **0.808** |
| Face Surprise Recognition (RT) | **0.806** |
| Face Delayed Matching (RT) | **0.718** |

**Supplementary Table 3:** **Composite scores for all participants**

| **Number** | **Participant** | **Laterality of lesion** | **Age** | **WORDS** | **OBJECTS** | **FACES** |
| --- | --- | --- | --- | --- | --- | --- |
| PL501 | Patient | L | 68 | 0,46 | 0,63 | -0,02 |
| PL502 | Patient | Bilat | 55 | 0,35 | 0,00 | -0,11 |
| PL503 | Patient | L | 87 | -3,54 | -2,92 | -2,33 |
| PL504 | Patient | R | 85 | -0,07 | -0,74 | -0,07 |
| PL505 | Patient | R | 69 | -1,15 | -1,44 | -2,66 |
| PL506 | Patient | L | 65 | 0,58 | 0,09 | 0,08 |
| PL507 | Patient | L | 70 | 0,01 | -1,13 | -1,07 |
| PL508 | Patient | L | 62 | -0,06 | 0,21 | 0,22 |
| PL510 | Patient | L | 67 | -0,06 | -0,51 | -0,65 |
| PL511 | Patient | L | 52 | 0,61 | 0,96 | 1,16 |
| PL513 | Patient | Bilat | 66 | -1,21 | -1,10 | -2,50 |
| PL514 | Patient | R | 71 | 0,05 | 0,00 | 0,58 |
| PL515 | Patient | L | 60 | -0,42 | 0,67 | 0,55 |
| PL516 | Patient | L | 65 | -0,35 | 0,14 | -0,63 |
| PL517 | Patient | R | 62 | 0,58 | 0,22 | 0,70 |
| PL518 | Patient | Bilat | 52 | 0,40 | -0,07 | -1,47 |
| PL519 | Patient | R | 52 | 0,17 | 0,50 | 0,60 |
| PL520 | Patient | R | 52 | -0,53 | -0,17 | -0,46 |
| PL521 | Patient | R | 62 | -0,12 | -0,53 | -2,23 |
| PL522 | Patient | R | 57 | 0,00 | -1,65 | -0,26 |
| PL523 | Patient | L | 57 | 0,50 | 0,28 | 0,83 |
| PL524 | Patient | L | 61 | 0,63 | 0,25 | 1,08 |
| PL525 | Patient | L | 80 | -0,70 | -0,46 | 0,17 |
| PL526 | Patient | Bilat | 65 | -3,96 | -3,58 | -3,54 |
| PL527 | Patient | L | 76 | 0,04 | -0,01 | -0,37 |
| PL528 | Patient | R | 84 | -0,18 | -1,44 | -1,02 |
| PL529 | Patient | L | 74 | -2,99 | -1,54 | -1,61 |
| PL530 | Patient | L | 64 | -0,28 | -0,22 | -0,21 |
| PL531 | Patient | L | 68 | -1,40 | -0,72 | -0,13 |
| PL533 | Patient | R | 70 | 0,18 | 0,34 | 0,12 |
| PL534 | Patient | L | 75 | 0,36 | 0,44 | 0,09 |
| PL535 | Patient | R | 71 | 0,45 | -0,33 | -0,41 |
| PL536 | Patient | R | 56 | 0,58 | 0,65 | 0,91 |
| PL537 | Patient | R | 27 | 0,13 | 0,33 | 0,31 |
| PL538 | Patient | L | 83 | -1,00 | -1,93 | -2,56 |
| PL539 | Patient | R | 34 | 0,47 | 0,57 | 0,65 |
| PL540 | Patient | R | 28 | 0,61 | 0,83 | 0,73 |
| PL541 | Patient | Bilat | 34 | -0,47 | -1,38 | -1,74 |
| PL543 | Patient | Bilat | 52 | 0,29 | 0,62 | 0,04 |
| PL544 | Patient | R | 55 | -0,07 | -0,28 | 0,55 |
| PL545 | Patient | Bilat | 62 | -0,38 | -0,94 | -0,53 |
| PM001 | Patient | L | 66 | -4,17 | -2,47 | -1,68 |
| PM002 | Patient | L | 51 | -1,28 | -1,49 | 0,45 |
| PM004 | Patient | L | 58 | -2,78 | -0,24 | 0,11 |
| PM006 | Patient | Bilat | 67 | 0,03 | -0,87 | -1,46 |
| PM007 | Patient | L | 63 | -1,53 | -1,19 | -2,31 |
| PM008 | Patient | L | 47 | 0,49 | 0,87 | 0,49 |
| PM009 | Patient | Bilat | 65 | -2,81 | -3,73 | -2,58 |
| PM010 | Patient | R | 52 | 0,50 | 0,84 | 0,72 |
| PM011 | Patient | L | 71 | -0,68 | -1,70 | -1,26 |
| PM012 | Patient | R | 46 | 0,44 | 0,83 | 0,92 |
| PM014 | Patient | L | 38 | 0,51 | 0,89 | 0,34 |
| PM015 | Patient | L | 44 | 0,17 | 0,68 | 0,71 |
| PM018 | Patient | L | 42 | -2,96 | -2,48 | -1,50 |
| PM019 | Patient | L | 70 | -0,19 | -0,46 | -0,14 |
| PM021 | Patient | L | 74 | 0,42 | 0,32 | 0,31 |
| PM022 | Patient | L | 57 | 0,47 | 0,52 | 0,86 |
| PM023 | Patient | L | 71 | 0,58 | 0,44 | 0,84 |
| PM024 | Patient | R | 66 | -0,14 | -0,93 | -1,49 |
| PM025 | Patient | R | 70 | 0,19 | -0,16 | -0,43 |
| PM026 | Patient | R | 62 | 0,16 | -1,27 | 0,04 |
| PM028 | Patient | L | 60 | 0,33 | 0,28 | 0,09 |
| PM030 | Patient | R | 50 | 0,42 | 0,43 | 0,71 |
| PM031 | Patient | R | 51 | 0,67 | 1,16 | 1,01 |
| CL801 | Control | - | 57 | 0,17 | -0,06 | -0,04 |
| CL803 | Control | - | 62 | 0,47 | 0,93 | 0,40 |
| CL804 | Control | - | 38 | 0,57 | 0,94 | 0,97 |
| CL805 | Control | - | 56 | 0,23 | 0,09 | 0,67 |
| CL806 | Control | - | 55 | 0,36 | 0,60 | 0,86 |
| CL809 | Control | - | 34 | 0,35 | 0,49 | 0,22 |
| CL810 | Control | - | 60 | 0,63 | 0,89 | 1,00 |
| CL811 | Control | - | 35 | 0,69 | 0,96 | 0,68 |
| CL813 | Control | - | 79 | 0,46 | 0,55 | 0,43 |
| CL814 | Control | - | 30 | 0,57 | 0,90 | 0,95 |
| CL815 | Control | - | 73 | 0,43 | 0,41 | -0,20 |
| CL816 | Control | - | 48 | 0,42 | 0,53 | 0,75 |
| CL817 | Control | - | 30 | 0,68 | 0,65 | 0,57 |
| CL818 | Control | - | 50 | 0,41 | 0,74 | 0,76 |
| CM301 | Control | - | 70 | 0,58 | 0,31 | 0,28 |
| CM303 | Control | - | 42 | 0,60 | 1,02 | 1,03 |
| CM304 | Control | - | 57 | 0,70 | 0,84 | 0,34 |
| CM305 | Control | - | 54 | 0,59 | 1,04 | 1,06 |
| CM306 | Control | - | 45 | 0,70 | 1,19 | 0,94 |
| CM307 | Control | - | 70 | 0,56 | 0,50 | 0,67 |
| CM308 | Control | - | 71 | 0,58 | 0,97 | 0,61 |
| CM309 | Control | - | 62 | 0,51 | 0,73 | 0,98 |
| CM310 | Control | - | 72 | 0,44 | 0,58 | 0,48 |
| CM311 | Control | - | 80 | 0,57 | 0,08 | -0,08 |
| CM312 | Control | - | 72 | 0,41 | -0,26 | 0,46 |
| CM313 | Control | - | 69 | 0,53 | 0,11 | 0,23 |
| CM314 | Control | - | 26 | 0,60 | 1,26 | 0,66 |
| CM315 | Control | - | 67 | 0,58 | 0,77 | 0,35 |
| CM316 | Control | - | 70 | 0,40 | 0,31 | 0,12 |
| CM317 | Control | - | 67 | 0,66 | 0,58 | 0,86 |
| CM318 | Control | - | 67 | 0,54 | 0,11 | -0,79 |
| CM319 | Control | - | 76 | 0,23 | -0,15 | -0,06 |
| CM320 | Control | - | 66 | 0,35 | 0,43 | -0,25 |
| CM321 | Control | - | 84 | 0,17 | -0,44 | 0,08 |
| CM322 | Control | - | 69 | 0,46 | 0,58 | 0,60 |
| CM323 | Control | - | 68 | 0,42 | 0,10 | 0,45 |
| CM325 | Control | - | 68 | 0,38 | 0,94 | 0,01 |
| CM326 | Control | - | 62 | 0,51 | 0,67 | 0,26 |
| CM327 | Control | - | 75 | 0,20 | -0,31 | -0,19 |
| CM328 | Control | - | 78 | 0,45 | 0,40 | 0,38 |
| CM329 | Control | - | 64 | 0,75 | 0,69 | 0,74 |
| CM330 | Control | - | 71 | 0,60 | 0,60 | 0,74 |
| CM331 | Control | - | 73 | 0,62 | 0,97 | 0,92 |
| CM332 | Control | - | 67 | 0,52 | 0,75 | 0,89 |
| CM333 | Control | - | 71 | 0,48 | 0,35 | 0,84 |
| CM334 | Control | - | 70 | 0,54 | 0,70 | 0,80 |

**Supplementary Table 4: whole brain multiple regression analysis on the constituent PCA ROIs**

|  |  | Words (beta) | Objects (beta) | Faces (beta) |
| --- | --- | --- | --- | --- |
| Model 1 | L ILF | -0.62 *** |  |  |
| Model 2 | L ILF | -0.50 *** |  |  |
|  | L Occipital | -0.38 *** |  |  |
| Model 3 | L ILF | -0.49 *** |  |  |
|  | L Occipital | -0.39 *** |  |  |
|  | R pITG | -0.22 ** |  |  |
| Model 1 | Total Lesion Volume |  | -0.63 *** |  |
| Model 2 | Total Lesion Volume |  | -0.57 *** |  |
|  | L ILF |  | -0.23 * |  |
| Model 1 | Total Lesion Volume |  |  | -0.57 *** |
| Model 2 | Total Lesion Volume |  |  | -0.52 *** |
|  | R aITG |  |  | -0.23 * |
| Model 3 | Total Lesion Volume |  |  | -0.43 *** |
|  | R aITG |  |  | -0.22 * |
|  | Unilateral vs. Bilateral stroke |  |  | -0.22 * |
| Model 4 | Total Lesion Volume |  |  | -0.61 *** |
|  | R aITG |  |  | -0.21 * |
|  | Unilateral vs. Bilateral stroke |  |  | -0.27 ** |
|  | R Lingual gyrus |  |  | 0.29 * |

*p<0.05, **p<0.01, ***p<0.001
